# Covert spatial selection in primate basal ganglia

**DOI:** 10.1101/282236

**Authors:** Fabrice Arcizet, Richard J. Krauzlis

## Abstract

The basal ganglia are important for action selection. They are also implicated in perceptual and cognitive functions that seem far removed from motor control. Here, we tested whether the role of the basal ganglia in selection extends to non-motor aspects of behavior by recording neuronal activity in the caudate nucleus while animals performed a covert spatial attention task. We found that caudate neurons strongly select the spatial location of the relevant stimulus throughout the task even in the absence of any overt action. This spatially selective activity was dependent on task and visual conditions, and could be dissociated from goal-directed actions. Caudate activity was also sufficient to correctly identify every epoch in the covert attention task. These results provide a novel perspective on mechanisms of attention by demonstrating that the basal ganglia are involved in spatial selection and tracking of behavioral states even in the absence of overt orienting movements.

## Introduction

The basal ganglia are known to govern behavior by disinhibiting desired actions and inhibiting undesired actions (Hikosaka et al. 2000). The basal ganglia have also been implicated in perceptual and cognitive functions (Pasupathy and Miller 2005), such as the encoding of object values (Kim and Hikosaka 2013) and action values (Seo et al. 2012), and signals related to visual decision-making (Ding and Gold 2010; 2012). Neurons in caudate nucleus, one of the major input structures of the basal ganglia, display a number of decision-related signals as monkeys formulate their eye movement perceptual report during a visual motion discrimination task (Ding and Gold 2010), consistent with a possible role for the basal ganglia in selective attention. Computational studies have provided a general framework that might account for these diverse functions, suggesting that the basal ganglia act as an integration center that plays a crucial role in representing the “belief state” needed to guide action selection (Samejima and Doya 2007). If this view of basal ganglia function is correct, then during tasks involving spatial attention one might expect to find neuronal signals in the caudate related to spatial selection and the internal encoding of belief states, even when no overt action or goal-directed movement is required.

Spatial selectivity in the caudate nucleus has been studied principally during tasks requiring a goal-directed movement, either with the eyes (Takikawa et al. 2002) or arms (Kermadi and Joseph 1995). In both cases, a subset of caudate neurons exhibits a degree of spatial selectivity as the monkey anticipates a movement instruction. Caudate neurons also show some spatial selectivity during the delay period preceding an action directed to the particular location (Hikosaka et al. 1989). In these paradigms, it is ambiguous whether the spatially selective activity is related to the visual location itself or to the spatial goal of the movement. In the antisaccade paradigm, which dissociates the visual target location from the movement endpoint, some caudate neurons have higher activity for antisaccades than for prosaccades (Ford and Everling 2009; Watanabe and Munoz 2010). However, even in this task, the instructional cue and movement endpoint are tightly linked, because antisaccades require a goal-directed eye movement to the location diametrically opposite the visual cue. Consequently, it is not known whether caudate activity can be spatially selective when animals attend covertly to a particular visual location that has no link to the endpoint of a goal-directed movement.

Another well documented contribution of the basal ganglia in the primate is the coordination of motor outputs by grouping individual movements into action “chunks” (Graybiel 1998) during sequences of eye movements (Seo et al. 2012) or arm movements (Miyachi et al. 2002). A recent study in rats found that striatal neurons were activated sequentially throughout the course of the delay period when animals had to wait before making a response, suggesting that sequence-related activity in the striatum might be a component of spatial working memory (Akhlaghpour et al. 2016). These studies raise the possibility that sequence-related activity in the primate striatum might also apply to the successive behavioral states that subjects pass through during the performance of covert spatial selection tasks in the absence of goal-directed or orienting movements, but this possibility has not yet been directly tested.

To test whether the primate striatum plays a more general role in spatial selection and in the internal representation of belief states in the absence of overt movements, we examined the activity of caudate neurons while monkeys performed a covert attention task. Animals were trained to covertly monitor a peripheral visual motion stimulus, and report when the direction of motion changed by releasing a joystick; unlike previous studies, the task required spatial selection but did not include movements toward a spatial goal. Our results demonstrate that the primate striatum is involved in covert spatial selection by showing that: 1) caudate neurons strongly discriminated the location of behaviorally relevant events, even though no goal-directed movement was involved, 2) this spatially selective activity required the presence of a distracter and often disappeared when only a single visual stimulus was presented indicating that the spatial selectivity was not only related to reward expectation and 3) the pattern of activity across the population of caudate neurons was sufficient to correctly identify the epochs in the attention task. By showing that caudate neurons are involved in spatial selection and tracking of behavioral states even in the absence of overt orienting movements, our results illustrate a possible common thread between the motor and cognitive functions of the basal ganglia.

## Material and Methods

### Animals

Data were collected from two adult monkeys (*Macaca mulatta*; Monkey R, 11 kg; monkey P, 14 kg). All procedures and animal care were approved by the National Eye Institute Animal Care and Use Committee and complied with the Public Health Service Policy on the humane care and use of laboratory animals. Under isoflurane and aseptic conditions, we surgically implanted plastic head-posts and recording chambers. In both animals, recording chambers (28 × 20 mm) were tilted laterally 35 degrees and aimed at the caudate head and body (20 mm anterior, 6 lateral).

### Experimental apparatus

The animals were seated in a primate chair (Crist Instrument Inc., Hagerstown, MD) with their head fixed inside a darkened booth. Animals were positioned 48 cm in front of a 100 Hz VIEWPixx display (VPixx Technologies, Inc., Saint-Bruno, QC, Canada), and experiments were controlled using a modified version of PLDAPS (Eastman and Huk 2012). Eye position was monitored using an EyeLink 1000 infrared eye-tracking system (SR Research Ltd., Ottowa, Ontario, Canada). Joystick release times (reaction times) were computed by detecting the onset of the step change in voltage from the joystick (CH Products, model HFX-10).

Neurons were recorded using tungsten in glass-coated electrodes with impedances of 1-2 MOhm (Alpha Omega Co., Inc., Alpharetta, GA). Electrode position was controlled with a stepping motor microdrive (NAN Instruments, Ltd, Nazaret Illit, Israel). The electrical signal was amplified, filtered and single-unit activity was recorded online using the Plexon MAP system spike sorting software (Plexon Inc., Dallas, Texas). Spike waveforms were analyzed again off-line to confirm that recordings were of single well-isolated neurons. We recorded neurons in the head and body of the left caudate nucleus for both animals with a range of AC+7 to AC-5 for monkey R and AC+5 to AC-2 for monkey P (AC: anterior commissure at AP20, Fig 1F). Neurons were considered to be in caudate nucleus according to their location (based on magnetic resonance imaging scans) and their low background activity at >10 mm below the dural surface. In this study, we recorded only from phasically active neurons (PANs), which we identified based on their low background activity compared to the tonic activity from the cholinergic interneurons (TANs). Single units with low or unstable firing rates across the session or with no task-related activity were excluded from the analysis.

**Fig 1.**
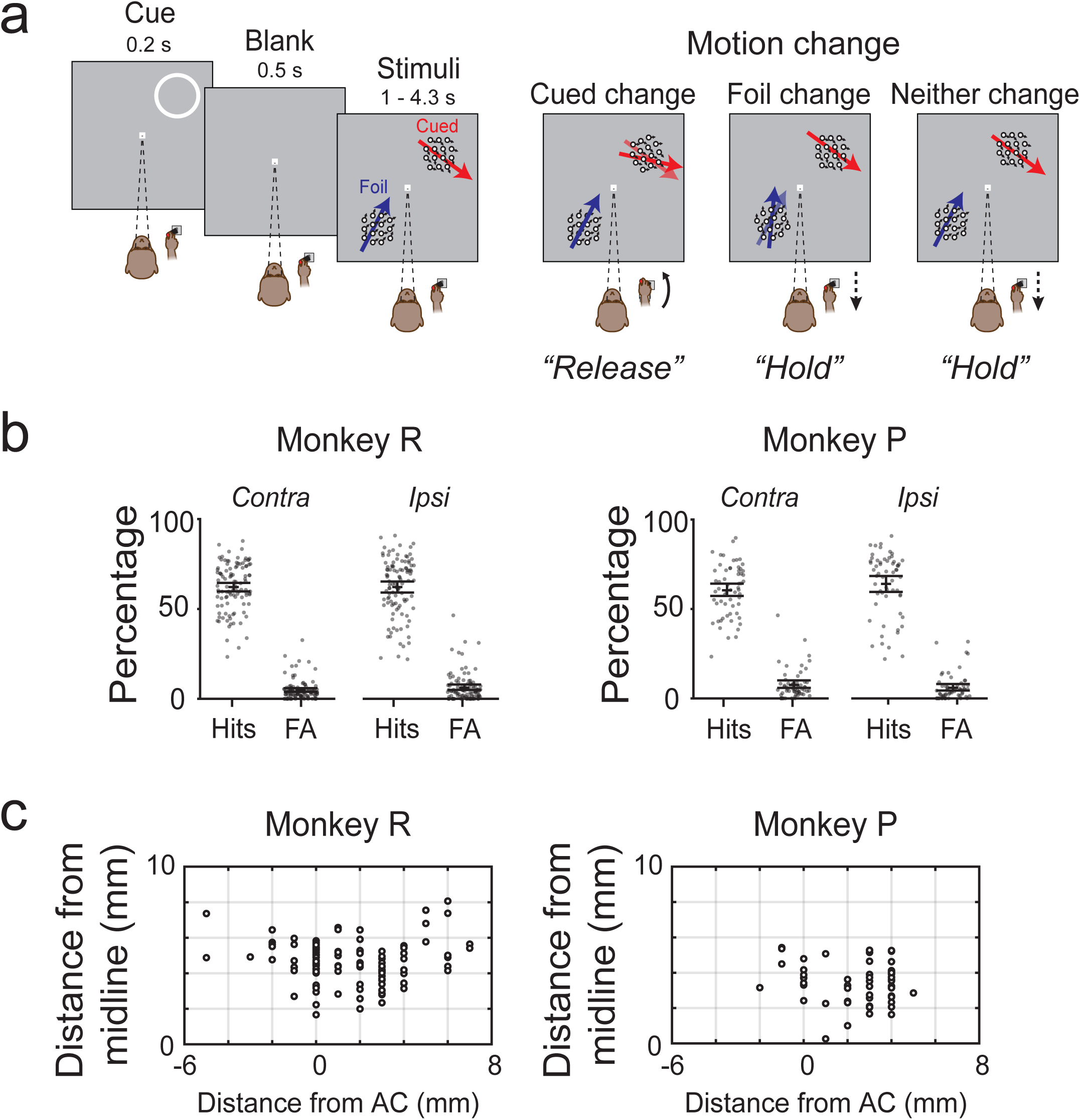
Change-detection task, behavioral performance and recording sites. (A) Task sequence in the covert attention task. While fixating a central spot and pressing down a joystick, a peripheral spatial cue (ring) was flashed for 0.2 s to indicate which part of the visual field the monkey should monitor. After a blank of 0.5 s, two motion patches were presented: one at the location previously cued (cued location) and the other one diametrically opposed (foil location). The monkey should detect when the motion direction changed at the cued location by releasing the joystick and otherwise keep holding the joystick down if the motion direction changed at the foil location or if no change occurred. (B) Behavioral performance in the task for both monkeys. Each dot represents the percentage of correct change trials (Hits) and false alarms (FA) for a single behavioral session (n=105 for monkey R, n=60 for monkey P), separately for trials with the cue contralateral or ipsilateral to the site of recording. Errors bars indicate 95 % confidence intervals of the mean. (C) Location of neuronal recording sites in the left caudate nucleus for both monkeys. Each dot represents one recording site. Positive values on x-axis indicate locations anterior to the anterior commissure (AC).

### Memory-guided saccade task and joystick task

Upon isolating spikes from a caudate neuron, we first tested neurons with a memory-guided saccade task in most cases (179/227, 78%). The monkey fixated a central spot for 0.5 s after which a spot stimulus was flashed for 0.15 s at one of 8 possible peripheral locations. The monkey maintained fixation until the fixation point turned off, at which point the monkey made a saccade to the memorized cued location within 0.5 s in order to receive a reward. We defined 4 different periods to test whether activity was modulated: visual [0-0.5s] and delay [0.5-1s] periods aligned on stimulus onset, saccade [-0.2:0.1s] and post-saccadic [-0.1:0.5s] periods aligned on saccade onset. 60% (108/179) of caudate neurons tested with memory-guided saccades were significantly modulated for at least one of these 4 temporal periods (Wilcoxon ranksum test, p<0.05 comparisons with baseline activity [-0.4:0s]), with the majority showing a preference for the contralateral side during the visual (55%) and saccade (57%) periods.

Because not all caudate neurons were modulated during memory-guided saccades, we also tested most neurons with a joystick task (190/227, 84%). The monkey held down the joystick and fixated a central spot in order for a peripheral white square stimulus to appear. On most trials (90%), after a random interval (1.5 – 3.5 s), the peripheral stimulus dimmed, and the monkey was rewarded for releasing the joystick within 1 second after the dimming. On 10% of trials, the peripheral stimulus did not dim, and the monkey was rewarded for continuing to hold down the joystick. The joystick task allowed us to check whether caudate neurons activities were modulated by joystick release, and by varying the location of the dimming stimulus, we could also assess preferences for visual stimulus location. 57% of caudate cells show significant activity during this joystick task at least during one of these 3 periods (visual [0:0.5s]; delay [0.5:1s] or joystick release [−0.2:0.1]) compared to a baseline period [−0.4:0s from stimulus onset]; most did not show any side preference (visual [29/14/57]; delay [23/12/65] and joystick release [27/21/52], for contralateral, ipsilateral or neither, respectively). A small proportion of caudate neurons (8%, 18/227) were not tested with memory-guided saccades or the joystick task, but only during the motion-direction change detection task.

### Motion-direction change detection task

All caudate neurons were tested using a motion-change detection task essentially identical to that described previously (Zenon and Krauzlis 2012), except we used a joystick rather than a button press. Briefly, the monkey started a trial by fixating a central white square and holding down a joystick for 0.25 s. A peripheral cue ring was then presented for 0.2 s to indicate which location in the visual field the animal should monitor. The cue was placed either at the neuron’s preferred location (as determined by memory-guided saccades or joystick task) or at the diagonally opposite location. The location of the ring was blocked for 68 successive trials. The cue ring was extinguished, and after 0.5 s when only the fixation point was still present, two motions patches (described below) were presented, one at the same location as the spatial cue, the other one (the foil) in the diagonally opposite location (Fig 1A). The direction of motion in the cued and foil patch were varied day to day, but always differed by 90 degrees.

We placed the motion stimuli at locations expected to evoke maximal activity for each neuron, based on the modulation observed during the memory-guided saccade and joystick task. The average eccentricity was 12 degrees (range: 10-13 degrees); in most cases, stimuli were placed on or near the horizontal meridian (<30 degrees). The visual motion stimuli were circular patches of moving dots, with the direction of motion of each dot drawn from a normal distribution with a mean value (defined as the patch motion direction) and a 16-degree standard deviation. The lifetime (10 frames, 100 ms), density (25 dots/deg^2^/s), and speed of the dots (15 deg/s) were held constant. The radius of the aperture varied between 3 to 3.75 degrees depending on the eccentricity of the patch; median value was 3.25 degrees. Luminance of the fixation dot and of each moving dot in the motion patches was 45 cd/m^2^. The background luminance of the monitor was 9.9 cd/m^2^.

The monkey was trained to release the joystick if the motion-direction changed in the patch at the previously cued location; otherwise he should keep holding the joystick down, on trials that had no motion-direction change or a motion-direction change in the foil patch. On each trial, a single motion-direction change could occur anytime 1.0 − 4.3 s after the onset of the motion stimuli. The proportions of trials with a motion-direction change at the cued location, foil location, or no change were 57%, 29 % and 14% respectively. The size of the motion-direction change was adjusted based on psychometric tests of each monkey to keep performance near threshold level (75% of performance), depending on visual field location and motion direction; the median direction changes were 28 and 26 degrees for monkeys R and P, respectively. Clockwise and counter-clockwise direction changes were equally likely and randomly chosen. After the motion-direction change, the stimuli remained on the screen for 1 second or until the animal released the joystick. Hits were defined as joystick releases that occurred within 1 second of a motion-direction change in the cued patch. False alarms were defined as incorrect joystick releases when the motion-direction change occurred at the foil location. Correct rejections were defined as successful non-releases when no change occurred at the cued location. Monkeys were rewarded with a small drop of liquid (apple juice mixed with water) at the end of each correctly performed trial (hits and correct rejections).

At the beginning of each block of trials, the monkeys performed the motion-change detection task with only one stimulus (single-patch condition) during the first 10-12 trials of each block. The single motion patch stimulus was located either at the cued location within the block (100% of the trials for Monkey R, 50% for Monkey P) or at the foil location (50 % of the trials for Monkey P). For Monkey P, the presentation of the single motion patch was preceded by the presentation of the spatial cue. Monkeys were rewarded for correct detection of motion change at the cued location and successful non-releases when no change occurred at the cued location (Monkey R and P) or when a change occurred at the foil location (Monkey P only). For analysis of single patch trials, we only used caudate activity during visual conditions that were common for both monkeys. Hit rates for single-patch conditions were 77.6% and 77.0% for monkeys R and P, respectively, confirming that the size of the motion-direction change was set near the threshold level.

Data from the motion-change detection task were collected in 105 recording sessions in monkey R and 60 recording sessions in monkey P. We obtained neuronal recordings (n=227 neurons) for an average of 200 trials per location of spatial cue; neurons recorded for fewer than 100 trials in the task were excluded from analysis. We defined contralateral trials as trials in which the spatial cue was presented on the side of the visual field contralateral to our recording site in the caudate. Monkeys were first trained on the task using their right hand (contralateral to the recording sites), which was used during all recordings except the 38 sessions (44 neurons) when the right and left hands were used in separate interleaved blocks.

### Neuronal PSTHs and cue-related modulation

For visualizing neuronal activity, we computed peri-stimulus time histograms (PSTHs) using non-overlapping bins of 0.02 s. To visualize neuronal activity across the population of caudate neurons, we computed normalized PSTHs by dividing the raw values from each time bin by the maximum firing rate (peak of each neuron’s PSTH). For analysis of neuronal activity, the firing rates within different temporal windows were computed from the trial-by-trial spike counts. To compare firing rates, we performed non-parametric statistical test as Wilcoxon signed rank (paired or not paired) or Kruskal-Wallis test with a significant threshold of p<0.05.

To quantify spatial cue-related modulation, we computed a standard attention cue modulation index defined as [R_contra_-R_ipsi_]/[R_contra_+R_ipsi_], where R_contra_ and R_ipsi_ is the mean activity on the contralateral trials and ipsilateral trials respectively. Mean activity was computed in different temporal windows: “pre-cue” [-0.25 to 0.02 s] before the spatial cue onset, “post-cue” [0.2 to 0.6 s] after the spatial cue onset, “visual” [0.1 to 0.5 s] after the motion stimuli onset, and “delay” [0.5 to 1 s] after the motion stimuli onset. We compared those mean activities with baseline activity [-0.95:-0.75 s] before the spatial cue onset (Wilcoxon signed rank p<0.05). This modulation index was also compute separately for trials in the single-patch condition.

### Analysis of response-choice activity

To analyze response-choice activity, we first identified neurons that showed significant changes in activity after the change in the visual motion stimulus. We aligned the data on the time of the motion-direction change and compared spike counts after the motion change [0.1 to 0.6 s] to those before the motion change [−0.5 to 0], and identified a subset of caudate neurons that had significantly higher activity after the motion change (80/227, 35%, Wilcoxon signed rank test, p<0.05). We restricted our analysis to this subset of caudate neurons. We aligned activity on the joystick release for hit responses (separately for contra and ipsi change trials), identified the time of peak activity by fitting a Gaussian function to the data from −0.5 to 0.5 s with respect to joystick release, and then measured the activity within a 0.3 s window centered on the peak. The spike counts from this 0.3 peak-centered window were then used for further analysis of response-choice activity.

We used a standard receiver operating characteristic (ROC) analysis (Britten et al. 1992) to determine the sensory and motor-related preferences of neurons during the response-choice epoch. For each neuron, we did three ROC-style analyses. The first analysis assessed how well response-choice activity discriminated the location of the visual motion change event. We divided correctly performed trials based on where the motion-direction change occurred (hits contralateral versus hits ipsilateral); for these two different types of trials, the action was the same (releasing the joystick) but the location of the visual event was different. The area under the ROC curve quantifies how well the location of the visual event could be discriminated based on the activity of each neuron, following a convention with values greater (less) than 0.5 indicating a preference for the contralateral (ipsilateral) side. The second analysis assessed how well response-choice activity discriminated the two behavioral outcomes (detect probability; hits versus misses); the sensory conditions were the same but the response choice was different (release versus hold). For this analysis, outcome values greater (less) than 0.5 indicating a preference for hits (misses). The third analysis assessed how well response-choice activity discriminated the presence or absence of the motion-change even in either motion patch (cued or foil). For this analysis, outcome values greater (less) than 0.5 indicating a preference for presence of the motion-change event (absence). To analyze data for trials without joystick releases, we aligned activity on the median reaction time separately for contralateral and ipsilateral conditions. Significance of ROC values was evaluated using bootstrapped (1000 iterations) 95% confidence intervals.

We also analyzed neuronal activity for three other cases in which the joystick was released: 1) false alarms, when the monkey incorrectly released the joystick for stimulus changes at the foil location, 2) joystick breaks, when the monkey incorrectly released the joystick when neither cued nor foil stimulus changed, and 3) joystick trial end, when the monkey appropriately released the joystick at the end of correct rejection trials to initiate the next trial. Only neurons with at least five occurrences for each type of these three cases were used for analysis. Spike counts for each neuron were measured from a 0.3 s window identical to that used to analyze response choice activity as described above.

### Classification of task epochs using a linear classifier

For each trial from every caudate neuron (n=227), we obtained spike counts in the motion-direction change task from 14 unique non-overlapping epochs, defined by different time periods during the trial (n=7) and whether the cue was ipsilateral or contralateral (n=2). The 7 time periods were: 1) “pre-cue”, a 0.23 s epoch starting .25 s before cue onset, 2) “post-cue”, a 0.4 s epoch starting 0.1 s after cue onset, 3) “visual”, a 0.4 s epoch starting 0.1 s after motion patch onset, 4) “delay”, a 0.5 epoch starting 0.5 s after motion patch onset, 5) “change contra”, a 0.5 s epoch starting 0.1 s after a motion-direction change in the contralateral patch, 6) “change ipsi”, a 0.5 s epoch starting 0.1 s after a motion-direction change in the ipsilateral patch, and 7) “change neither”, a 0.5 s epoch matched in time to the two preceding change epochs.

We first used the “svmtrain” and “svmclassify” functions in Matlab (version R2015b, The Mathworks, Natick, MA) to train and test a linear classifier for each of the 14 epochs defined above. For each classifier, we randomly drew (with replacement) a single-trial spike count from each neuron from the corresponding epoch to generate a single-trial feature set (n=227 neurons), and then repeated this procedure multiple times (n=150) to make the full data set for that classifier. 120/150 trials were used to train the classifier, and the remaining 30 trials were held in reserve to test and cross-validate the performance of the classifier. Each of the 14 classifiers was also tested with the reserve data from the other 13 classifiers to generate a confusion matrix (i.e., epochs that might be identified by more than one classifier). This procedure was then repeated 1000 times, and the 5^th^ percentile from the distribution of outcomes was compared to chance performance to identify significant results (reported as medians).

In a second analysis, we followed the same procedure using data from the epochs related to response choice (epochs 5-7 defined above, for contralateral and ipsilateral cue conditions), subdivided based on trial outcome (hit, correct reject, miss, false alarm: uncued change, and joystick breaks: no change). We trained classifiers for the 4 possible correct trial outcomes, and then tested these 4 classifiers on all 10 possible trial outcomes. A bootstrap procedure (1000 repeats) was used again to assess significance.

## Results

We recorded the single-unit activity from 227 caudate neurons in 2 monkeys (153 for monkey R and 74 for monkey P) trained to perform a motion-change detection task (Fig 1A). The neurons were identified as phasically active neurons (Hikosaka et al. 1989; Apicella et al. 1992) based on their low background activity and were located in the head and body of the caudate nucleus within 7 mm of the anterior commissure (Fig 1C). The neuronal data were qualitatively similar across the two animals and have been pooled for simplicity.

Before describing the neuronal data, here we briefly characterize the monkeys’ performance in the attention task. As described in detail in Materials and Methods, “hits” were defined as correct releases of the joystick when monkeys detected a change in the direction of motion at the previously cued location, and “false alarms” were defined as incorrect releases of the joystick when the motion-direction change occurred at the uncued foil location (Fig 1A). The task invoked mechanisms of spatial attention, because the amplitudes of the motion changes were placed near each monkey’s psychophysical threshold and the task required ignoring irrelevant changes in the direction of motion at the uncued foil location. Both monkeys performed the task reliably across a total of 165 recording sessions (Fig 1B). The hit rates of both monkeys were 60-64% (monkey R: 62.2% contra, 62.2 % ipsi; monkey P: 60.4 % contra, 64.0% ipsi) with only minor differences in hit rates between the two sides (R: p=0.996, P: p=0.070, Wilcoxon sign rank test). The false alarm rates for foil changes were low (R: 4.9% contra, 6.1% ipsi; P: 7.5% contra, 5.9% ipsi) and not different between the two sides (R: p=0.196; P: p=0.124, Wilcoxon sign rank test). Conversely, correct reject rates (i.e., non-releases of the joystick when neither the cue or foil stimulus changed) were 71-79% (monkey R: 71% contra, 74% ipsi; monkey P: 79% contra, 79% ipsi). The two monkeys showed small (less than 20 msec) but significant differences in mean joystick reaction time for stimulus changes on the two sides (monkey R: 0.579 sec contra, 0.588 sec ipsi, p=0.007; monkey P: 0.498 sec contra, 0.516 sec ipsi, p=0.005; Wilcoxon rank sum test). Analysis of the voltage traces used to measure the position of the joystick indicated that the movements used to release the joystick were nearly identical in both monkeys across these conditions. The absence of strong spatial biases in the behavior is noteworthy, because our sample of caudate neurons showed patterns of activity during the performance of the attention task that often depended on whether the cue was contralateral or ipsilateral to the recording site.

### Caudate activity related to the spatial cue and motion stimulus

Caudate neurons showed several distinctive patterns of activity during the early epochs of the attention task, when the spatial cue and the motion patches were presented. To illustrate the range of activity patterns, we show the time course of spike counts (peri-stimulus time histograms) from four example caudate neurons, sorted by whether the spatial cue was presented contralateral (orange) or ipsilateral (blue) to the recording site (Fig 2A).

**Fig 2.**
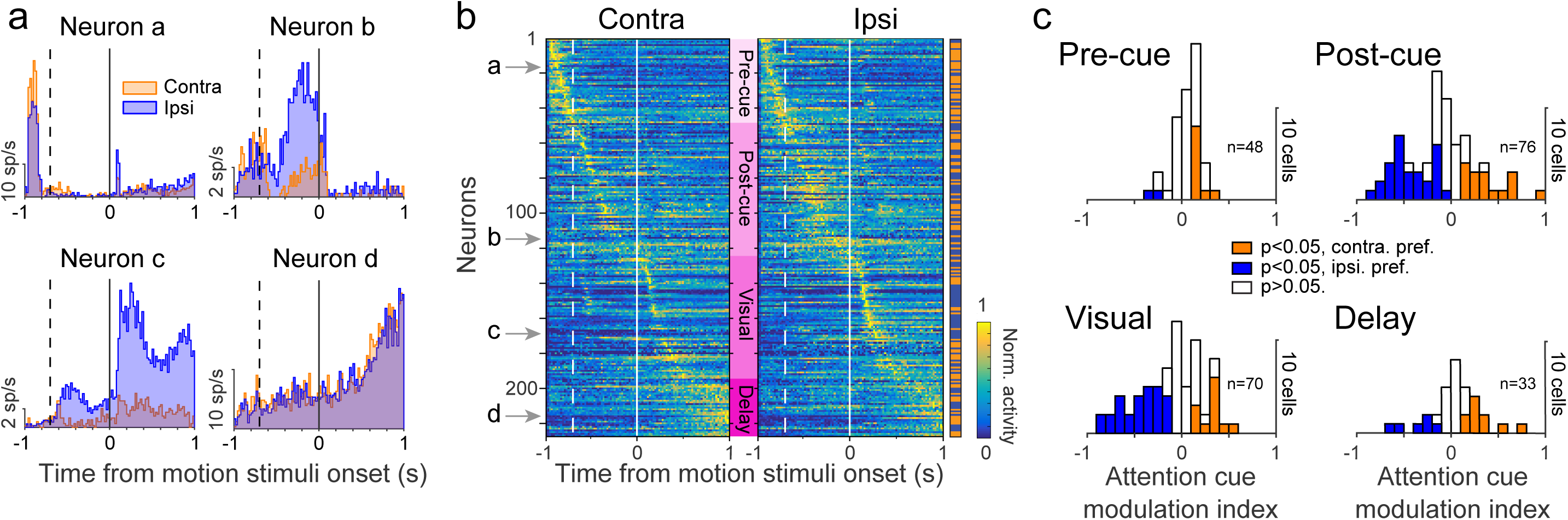
Caudate neuronal activity and cue-related modulation during the change-detection task. (A) Example caudate neurons. Activity of four different caudate neurons (a, b, c, and d) aligned on the onset of the motion patches (solid vertical line) when the cue was contralateral (orange) or ipsilateral (blue) with respect to the recording site. The dashed vertical lines indicate when the spatial cue was presented. (B) Normalized activity for our complete population of caudate neurons (n=227). Each row represents the normalized activity of a single neuron for contralateral presentation of the spatial cue (left) or ipsilateral presentation (right) aligned on motion stimuli onset. Neurons were ranked according to the time of their peak activity across both contralateral and ipsilateral cue conditions. Colored sidebar on the right indicates whether each neuron had maximal activity for contralateral (orange) or ipsilateral (blue). Solid white lines indicate onset of motion stimuli, dashed lines spatial cue onset. Neurons were grouped according to the timing of the peak activity (labels in pink): before the spatial cue onset (Pre-cue), between the spatial cue and the motion stimuli onset (Post-cue), after the motion stimuli onset (Visual), and during the delay period prior to motion-direction change (Delay). (C) Spatial cue modulation in caudate nucleus. The histograms display the distribution of attention cue modulation index values for the periods for each group of neurons.

The activity of many caudate neurons was related to the location and timing of the spatial cue. Some of these neurons exhibited phasic activity that preceded the appearance of the spatial cue (Neuron a, Fig 2A), suggesting the presence of an anticipatory signal made possible by the 68-trial blocking of spatial cue conditions and the fixed temporal period (0.25 s) before the appearance of the spatial cue. The pre-cue activity of this neuron was slightly but significantly larger when the cue was contralateral than when it was ipsilateral (p=0.017, Wilcoxon ranksum test, period [−0.250 to −0.020 before cue onset],18.5 sp/s on average for contra trials vs 14.4 sp/s for ipsi trials). Other neurons showed phasic activity after, and presumably evoked by, the spatial cue (Neuron b). The post-cue activity for this neuron was much larger when the cue was presented on the ipsilateral side (Wilcoxon ranksum test, period [0.1 to 0.5 s after cue onset], p<0.001, 0.5 sp/s for contra trials vs 4.1 sp/s for ipsi trials).

For other caudate neurons, activity was mostly related to the presentation of the motion stimuli and the delay period of the attention task. Some neurons exhibited large phasic responses to the onset of the motion stimuli, followed by activity that extended into the delay period of the task (Neuron c, Fig 2A), with a strong preference based on the location of the spatial cue that emerged shortly after cue onset; this side preference nearly eclipsed the visual phasic response in the non-preferred condition, even though the visual stimuli were identical across the two cue conditions. The delay period activity also varied across caudate neurons. Some showed a distinctive ramp-like pattern toward the end of the delay period without any side preference (Neuron d).

To visualize the activity patterns across our population of caudate neurons (n=227) during the early phases of the task, we normalized each neuron’s spike counts and rank-ordered all of the neurons based on the times of their peak activity (see Materials and Methods). The iconic representation of these results (Fig 2B), aligned on motion stimulus onset separately for contralateral and ipsilateral cue conditions, illustrates that the sample neurons in Fig 2A were exemplars of features present across the population. Specifically, based on the timing of peak activity, we classified neurons into four different groups (indicated by labels in pink gradient). The first group of neurons (Pre-cue, light pink, n=48) showed peaks of activity preceding the appearance of the spatial cue (like Neuron a). Neurons in this group tended to exhibit phasic activity for both contralateral and ipsilateral cue conditions, with slightly higher activity for contralateral (as indicated by both the higher normalized activity for the contra plot and the larger proportion of orange horizontal tics in the side bar of Fig 2B).

The second group of neurons (Post-cue, n=76) increased their activity after the presentation of the spatial cue (like Neuron b). These “post-cue” neurons tended to show higher activity in the ipsilateral cue condition (as indicated by the higher normalized activity and the larger proportion of blue tics in the side bar of Fig 2B). The timing of this activity was distributed across the post-cue epoch, including just after cue onset (neurons #50-60), after cue offset (neurons #80-100), and just before motion stimuli onset (neurons #115-120).

The third group of neurons (Visual, n=70) had responses that appeared to be evoked by the onset of the motion stimuli (like example Neuron c). These “visual” neurons sometimes also exhibited cue-related activity before motion stimuli onset, and lower sustained activity into the delay period. The phasic visual response showed a preference for the ipsilateral cue condition (neurons #140-170), whereas the preferences during the delay-period were more equally split.

For the last group of neurons (Delay, n=33) the peak of activity occurred well after the presentation of the motion stimuli and into the delay period (0.5 s and longer after motion stimuli onset). These “delay” neurons tended to show a ramping pattern of activity (like example Neuron d), similar to that described previously by Ding and Gold (2010), and had a slight preference for the contralateral cue condition.

### Modulation of caudate neurons by contralateral and ipsilateral cues

To quantify the cue-related modulation, we computed a modulation index for spike counts within different temporal periods (Fig 2C, see Material and Methods) and found clear distinctions between the groups of caudate neurons. The neurons with prominent pre-cue activity (n=48) were weakly modulated by the spatial cue condition (Fig 2C). Only 25% (12/48) of this group showed significantly different activity based on cue condition, with most preferring contralateral (10/12), and the amplitude of this modulation was small (median value of modulation indices of 0.16 for the 10 cells that preferred contralateral side, orange filled bars). In contrast, the post-cue and visual neurons were strongly modulated by spatial cues. Most of the post-cue neurons (45/76, 58%) displayed a significant effect of cue condition, with almost 2/3 showing a preference for ipsilateral (29/45, 63%). Similarly, a majority of visual neurons (40/70, 57%) showed significant cueing effects, again mostly in favor of the ipsilateral cue condition (28/40, 70%). The amplitude of the cueing effects for post-cue and visual neurons was large – the median cue modulation index was − 0.49 and −0.43 (post-cue and visual, respectively) for the ipsilateral condition, and 0.32 and 0.32 for the contralateral condition. Finally, neurons defined by their delay period activity were also modulated by spatial cues; half of the neurons (16/33) showed a significant effect, with a slight preference for the contralateral side (10/16), but the size of the modulation was smaller (median: 0.19 and −0.33 for contra and ipsi, respectively).

### Dependence of spatial selectivity on the visual configuration

We unexpectedly found that the spatial cue modulation was dependent on the presence of a visual distracter during the covert attention task. For visual neurons (n=70), we compared activity during the standard two-patch version of the attention task with activity during a simpler single-patch version that omitted the distracter. During this single-patch task, a single motion patch was presented at the contralateral or ipsilateral location, and the animal was again rewarded for releasing the joystick if the motion direction in the single patch at the cued location changed.

This comparison revealed that the visual activity of many caudate responses was selective for visual conditions that included the second distracter patch. For example, Neuron#1 in Fig 3A showed a strong preference during the two-patch attention task for ipsilateral placement of the cued stimulus (blue vs orange, top left quadrant). This preference completely disappeared during the single-patch condition (lower left quadrant), demonstrating that the spatial selectivity of this neuron was specific to visual conditions in which an ipsilateral cued stimulus was accompanied by a contralateral distracter. Some caudate neurons were like Neuron #1 and displayed spatial selectivity during the two-patch version of the covert attention task, but lost their side preference during the single-patch condition (green dots in Fig 3B, n=27/70, significant AMI for two-patch but not single-patch); a majority of these neurons (20/27) had a preference for the ipsilateral side (i.e., AMI < 0 for the two-patch condition).

**Fig 3.**
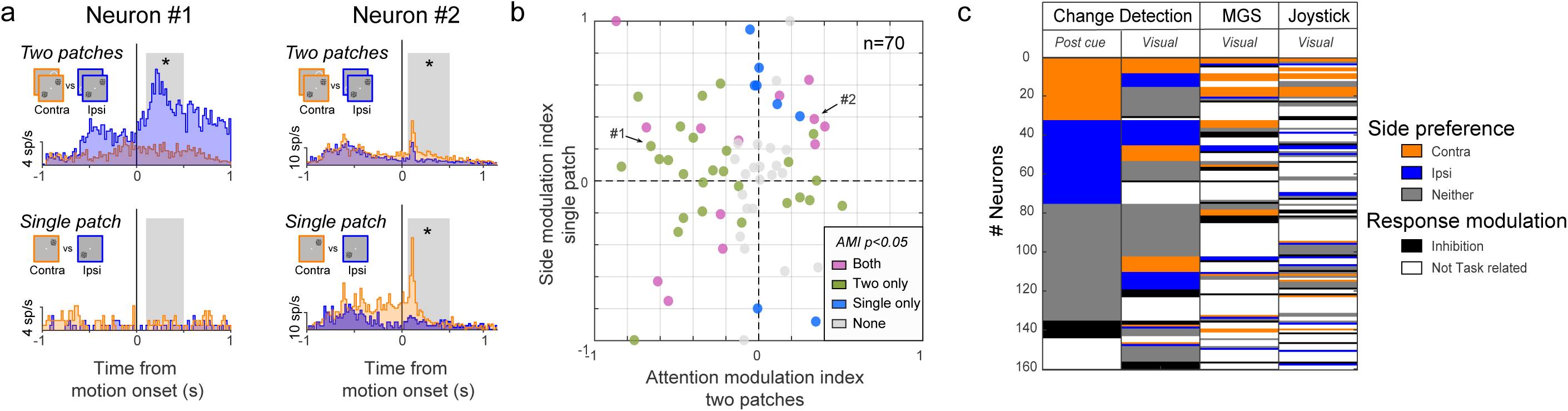
Dependence of caudate visual activity on the presence of distracters. (A) Firing rates for two sample caudate neurons aligned on presentation of visual stimuli for the two-patch condition (top row) and single-patch condition (bottom row). The orange/blue code indicates respectively the spatial cue location or the single patch location for contra/ipsi side. Gray boxes demarcate the 0.4 s time period used for computing the Attention Modulation Index (AMI) for the two-patch condition and Side Modulation Index for the single-patch condition (SMI). Asterisks indicate significant values for AMI/SMI (Wilcoxon rank, p<0.05). (B) Scatter plot of AMI and SMI values computed for each caudate neuron with visual activity (n=70). AMI values on x-axis greater (less) than 0 indicate preference for contralateral (ipsilateral) hits in the two-patch condition. SMI values on y-axis greater (less) than 0 indicate preference for contralateral (ipsilateral) for single-patch condition. Each dot represents one caudate neuron from “visual” subpopulation (Fig 2B, n=70). Color indicates the neuron’s group assignment based on the AMI/SMI values: AMI for two patches different for one side (green), both different from chance (purple), SMI for single patch different for one side (blue) and neither different from chance (gray). (C) Side preference for the motion change-detection task, memory guided saccade task (MGS) and joystick task in the population of 160 caudate neurons tested with the 3 tasks. Each row represents a single neuron across different visual epochs: post-cue and visual for change detection task, visual periods for MGS and joystick mapping task. Neurons were sorted according their side preference for post-cue period. Colors indicate side preference (orange for contra, blue for ipsi, gray for neither) when the activity was significantly greater than the baseline (Wilcoxon rank-sum test, p<0.05), when no significant activity was reported (Wilcoxon rank-sum test, p≥0.05; white) or when activity was significantly lower than the baseline (Wilcoxon rank-sum test, p<0.05; black).

Less common were neurons like Neuron #2 (Fig 3A), which exhibited a side preference during both single-patch and two-patch conditions (violet dots, n=13); most of these neurons (9/13) retained a consistent side preference across visual conditions like Neuron #2. The 4 remaining neurons preferred the ipsilateral side in the two-patch condition but preferred the contralateral side in the single-patch condition. Finally, some neurons (gray dots, n=22/70) did not show a side preference during either the single-patch or two-patch condition.

The visual activity of many caudate neurons was selective to the visual configuration during the covert attention task and could not be predicted by either the memory guided saccade (MGS) or joystick mapping tasks. In a population of 160 neurons tested in all three tasks, we found that most caudate neurons did not show the same spatial selectivity (Fig 3C) across the different tasks. For the neurons that preferred the contralateral side during the post-cue period of the attention task, (left column, orange rows, n=32/160), only 10/32 showed the same spatial selectivity for MGS and joystick task during the visual periods and only 2/43 neurons showed congruent selectivity for the ipsilateral side.

In summary, we observed cue-related modulation across all of the early epochs of the attention task, indicating that these cueing effects were not related to the delivery of the reward or to behavioral outcomes at the end of the trial. The effects were strongest in the post-cue and visual epochs, and larger for ipsilateral than contralateral spatial cues. The visual cue-related modulation for many caudate neurons depended on the visual configuration – it was specific to the change detection paradigm, required the presence of a visual distracter, and disappeared when only a single visual stimulus was presented at the preferred location.

### Activity of caudate neurons related to response choice

As might be expected from previous results implicating the caudate nucleus in movement sequencing and procedural learning, a subset of our caudate neurons was modulated during the joystick response choice. However, even among these neurons, neuronal activity was not simply movement-related, but exhibited unexpected selectivity for the visual and task conditions.

Among the subset of neurons with activity modulated during the response choice (n=80, defined in Material and Methods), we found that response-choice activity could be driven by sensory signals (i.e., where the motion change happened), motor-related signals (i.e., whether or not the animal released the joystick), or combinations of the two. For example, the response-choice activity of Neuron #1 in Fig 4A combined a preference for contralateral over ipsilateral motion-direction changes (top) with a preference for hits over misses (bottom). To quantify these sensory and choice-related signals, we used ROC analyses to measure: 1) the neuronal motion-change selectivity, by comparing spike counts on contralateral hits to ipsilateral hits (AROC sensory), and 2) detect probability, by comparing spike counts on trials with hits versus misses (AROC motor). For Neuron #1, both ROC analyses were both significantly greater than chance (AROC of 0.88 and 0.6, sensory and motor, respectively), confirming that the response-choice activity of this neuron combined a preference for hits with a preference for stimulus changes in the contralateral visual field. In contrast, Neuron #2 preferred hits over misses (AROC: 0.71) but had no preference between contralateral and ipsilateral change-event locations (AROC: 0.5), suggesting that the response-choice activity of this neuron was predominantly related to the joystick release. Similar mixed dependencies were observed across the caudate neurons with response-choice activity. Some caudate neurons were like Neuron #1 and combined a preference for change-event location with a preference for hits over misses (green dots in Fig 4B, n=32/80, both AROC were significantly different from chance level 0.5); most (23/32) of these neurons preferred contralateral change-events (AROC > 0.5) with an almost exclusive preference for hits (31/32). Almost as common were neurons like Neuron #2, which signaled the motor choice without a preference for change-event location (red dots, n=30). Finally, a smaller number of neurons (blue dots, n=13) exhibited response-choice activity that discriminated the location of the motion change (and generally preferred contralateral) but did not show any difference between hits and misses. Despite these distinctions, caudate neurons did not form exclusive categories based on their response-choice activity, but showed a continuum of mixed preferences for change-event location and motor choice, as illustrated by the broad scatter of data points in Fig 4B.

**Fig 4.**
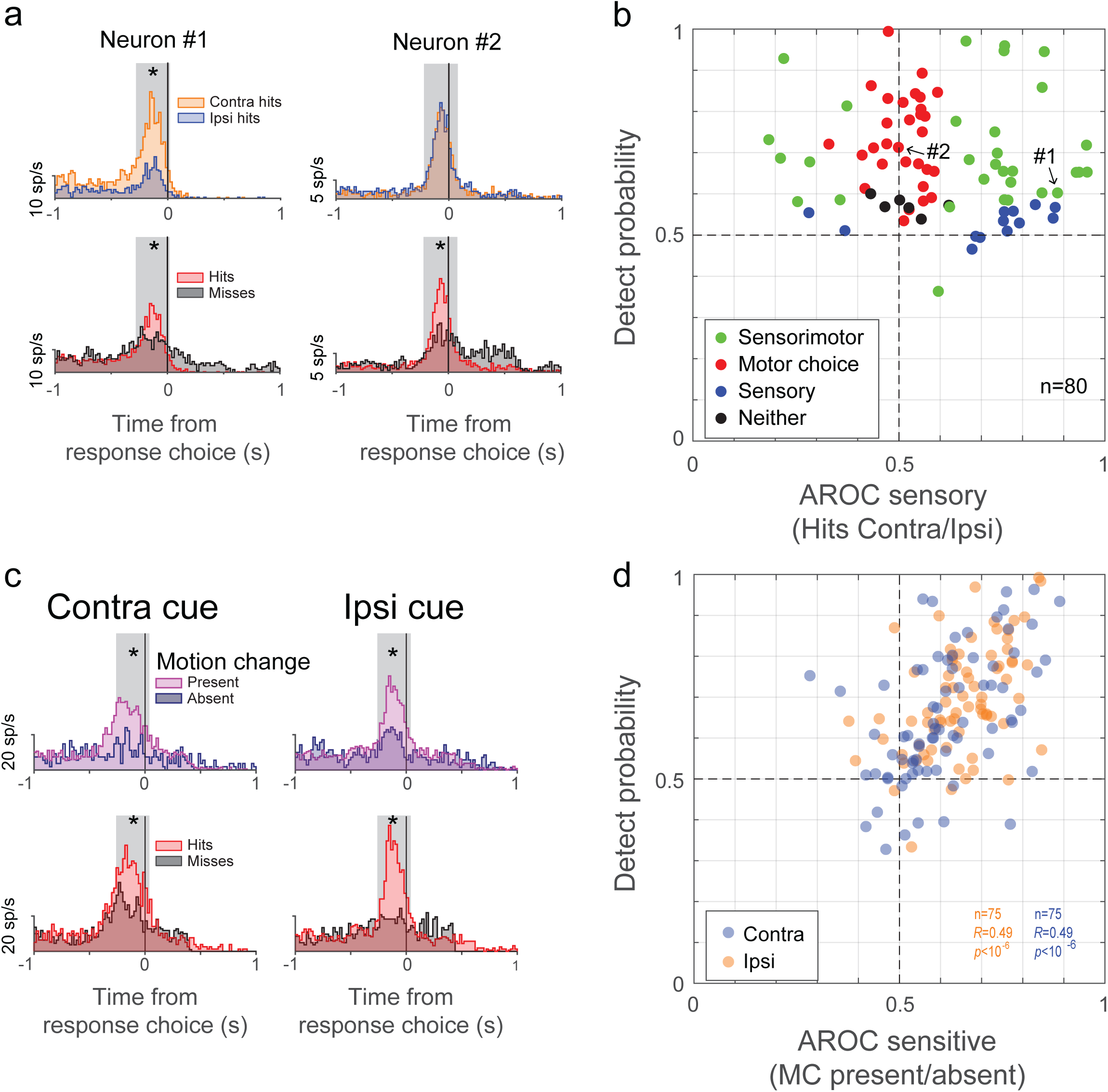
Response-choice activity in caudate nucleus. (A) Firing rates for two sample caudate neurons (#1 and #2) aligned on the joystick release. Upper row shows the response-choice activity for contralateral hits (orange) and ipsilateral hits (blue); lower row shows activity for hits (red) and misses (gray) pooled across stimulus locations. Grey boxes demarcate the 0.3 s time period used for the ROC analysis. Top asterisk indicates that the AROC value was significantly different from chance level (0.5). For miss trials, which did not contain joystick releases, data for each neuron were aligned on the median reaction time during the recording session. (B) Scatter plot of detect probabilities and sensory ROC values computed for each caudate neuron that showed response-choice activity. AROC values on x-axis greater (less) than 0.5 indicate preference for contralateral (ipsilateral) hits. Each dot represents one caudate neuron (n=80). Color indicates the neuron’s group assignment based on the AROC values: hits vs. misses AROC different from chance (“motor choice”, red), contralateral hits vs. ipsilateral hits different from chance (“sensory”, blue), both different from chance (“sensorimotor”, green), neither different from chance (black). (C) Firing rate for one caudate example aligned on the joystick release. Upper row shows the response-choice activity in the presence (pink) or absence (blue) of the motion change; lower row shows activity for hits (red) and misses (black). For the motion change present trials (pink), the change event could happen at either location (cued or foil). Left column shows responses for the contra trials, right for the ipsi trials. Grey boxes demarcate the 0.3 s time period used for the ROC analysis. Top asterisk indicates that the AROC value is significantly different from chance level (0.5). For miss trials and some motion-change absent trials, which did not contain joystick releases, data for each neuron were aligned on the median reaction time during the recording session. (D) Scatter plot of the detect probabilities as a function of the neural sensitivity (AROC “sensitive”) for the caudate neurons with response-choice activity. Only neurons with at least five occurrences for each type of conditions were used for analysis (n=75/80). The color code indicates the location of the spatial cue; contra (orange) and ipsi (blue). AROC values on x-axis greater (less) than 0.5 indicate preference for presence (absence) of the motion change; AROC values on y-axis greater (less) than 0.5 indicate preference for hits (misses).

We next tested the modulation of caudate responses to the presence or absence of the change event. Fig 4C shows an example of a typical caudate neuron whose activity was significantly modulated by the presence or absence of the change-event and also for hits versus misses, independently of the cue location (contra/ipsi). We measured the correlation between neuronal sensitivity and detect probability across our population of neurons, separately for contralateral and ipsilateral cue conditions. Neuronal motion-change sensitivity was significantly and positively correlated with the detect probability for both contralateral and ipsilateral cueing conditions (Fig 4D, contra trials, *R* = 0.49, P < 10^−6^; ipsi trials, *R* = 0.49, P < 10^−6^), indicating that caudate neurons with greater sensitivity to the visual event were also more strongly predictive of the response choice.

Together, these results show that the response-choice activity of caudate neurons was selective for the spatial location of the relevant stimulus event, and that this activity was correlated with the animal’s behavioral choice during the attention task.

### Dependence of response-choice activity on attention task conditions

The phasic response-choice activity was also dependent on the task condition in which the joystick was released. We considered three other situations during the covert attention task when the animal released the joystick, in addition to the case of “hits” analyzed in Fig 4. First, we analyzed activity during “false alarms”, when the animals incorrectly released the joystick for a motion change that occurred at the foil location. Second, we analyzed “joystick breaks”, when the animals released the joystick but there was no motion change event at either stimulus location. Third, we analyzed activity during “joystick releases” at the end of correct reject trials, when the animal was obliged to release the joystick in order to end the trial.

Mean activity for false alarms was not different from that for hits for most of the neurons (Fig 5A; 10/14, 71% contra; 12/14, 86% for ipsi). Similarly, mean activity for joystick breaks was equivalent to activity for hits for most of the neurons (Fig 5B; 34/46, 74% contra; 26/31, 84% for ipsi). In contrast, mean activity for hits was significantly larger than mean activity for joystick release at the end of correct-reject trials (Fig 5C, Wilcoxon signed rank, p<0.001 for contra, p<0.001 for ipsi). These results indicate that the response-choice activity was specific to joystick releases associated with the choice about the visual motion stimulus.

**Fig 5.**
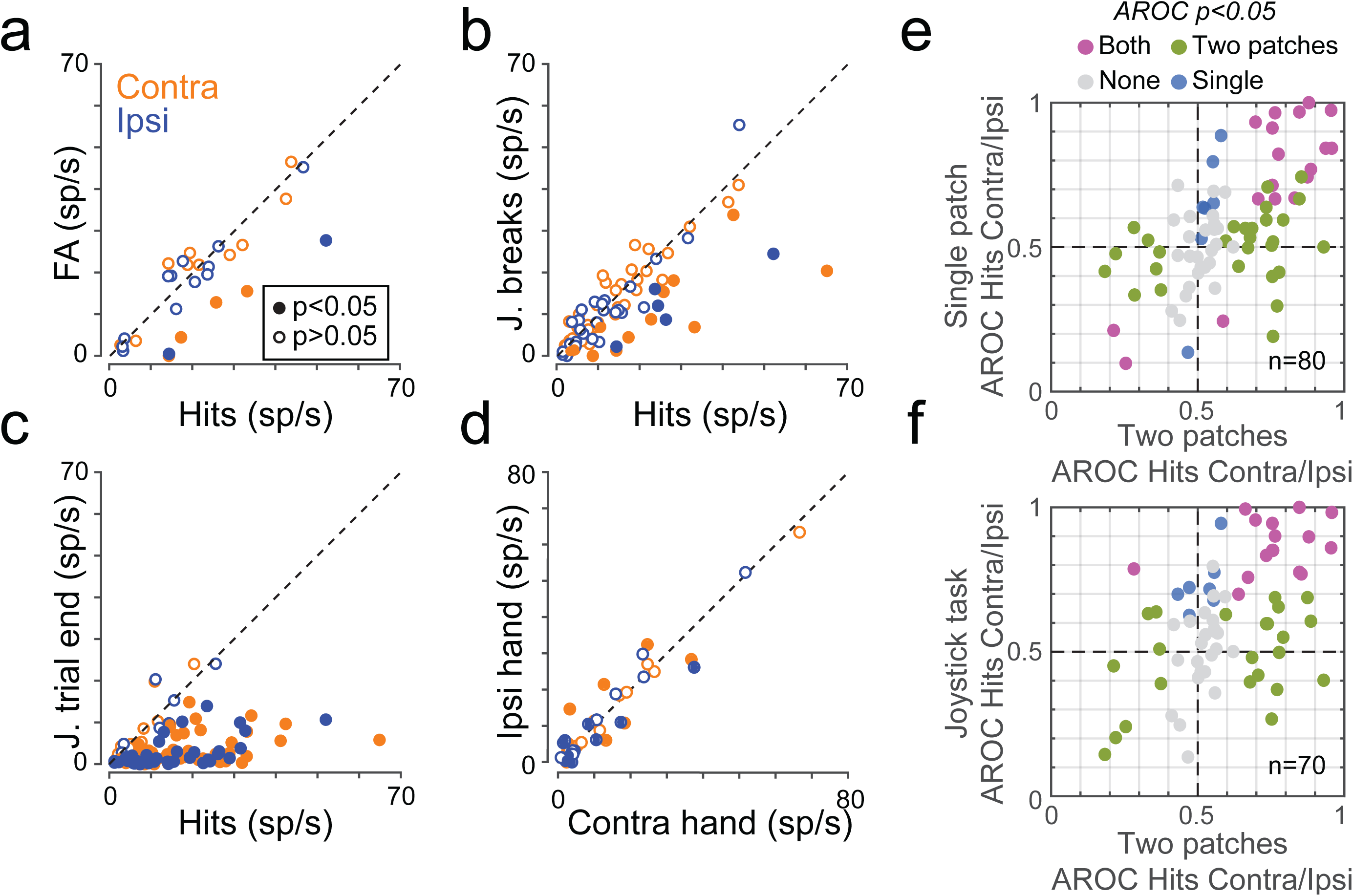
Error trials and task influence. (A, B, C). Scatter plots of mean caudate activity during false alarms (A), joystick breaks (B), and joystick trial end (C) as a function of mean activity to hits for contralateral trials (orange) and ipsilateral trials (blue). Filled dots (orange or blue) indicate when mean activities were significantly different (Wilcoxon rank sum test, p<0.05). Dashed lines represent identity lines. Joystick breaks are defined as trials when the animals released the joystick but there was no motion change event at either stimulus location and “joystick releases” at the end of correct reject trials, when the animal was obliged to release the joystick in order to end the trial. (D) Scatter plot of mean responses for contra and ipsi hits (orange and blue) computed during the 0.3 s time period when animals used either their ipsi hand (y-axis) or contra hand (x-axis) with the joystick. Same convention as A, B and C. E) Scatter plot for each caudate neuron (n=80) with response-choice activity comparing AROC values computed for single-patch and two-patch conditions. AROC values on x-axis greater (less) than 0.5 indicate preference for contralateral (ipsilateral) hits for two-patch condition; AROC values on y-axis greater (less) than 0.5 indicate preference for contralateral (ipsilateral) hits for single-patch condition. Color indicates the neuron’s group assignment based on the AROC values: contralateral hits vs. ipsilateral hits different from chance for two patches only (green), contralateral hits vs. ipsilateral hits different from chance for single patch only (blue), contralateral hits vs. ipsilateral hits different from chance for both conditions (purple), neither different from chance (gray). (F) Scatter plot of AROC values computed for each caudate neuron tested with the dimming joystick task (n=70) with response-choice activity for two-patch and dimming joystick conditions. AROC values on y-axis greater (less) than 0.5 indicate preference for contralateral (ipsilateral) hits for dimming joystick condition.

As an additional test of the specificity of this response-choice activity, for a subset of caudate neurons we tested whether the hand used to release the joystick made a difference. We defined contra and ipsi hand relative to the recording site of the neurons, as we did for spatial cue location. We recorded a total of 44 neurons, in 38 behavioral sessions (n=25 for monkey R and n=13 for monkey P). Behavioral performance was not different when animals used their contralateral hand or the ipsilateral hand with only minor differences in hit rates between the two hands (R: 65.2% ipsi hand, 66.3% contra hand, p=0.777, Wilcoxon signed-rank test; P: 57.7% ipsi hand, 65.4% contra hand, p=0.002). Among the population of neurons that showed choice-related phasic activity (n=20), there was no preference for one particular hand. Indeed, mean responses for correct responses to motion changes on the contralateral side (orange dots) or ipsilateral side (blue dots) did not depend on which hand was used to release the joystick (Wilcoxon rank, p=0.681 contra change, p=0.575 ipsi change, Fig 5D).

In summary, the features of the phasic activity related to joystick release support the use of the term “response-choice” – this activity was specific to joystick releases associated with choices about the visual motion stimulus, but was largely unaffected by which hand was used.

However, the response-choice activity was strongly affected by the visual and task conditions (Fig 5E,F). We compared caudate neuronal activity for joystick releases during three different conditions: 1) the standard two-patch attention task, 2) the single-patch version of the attention task introduced earlier, and 3) a joystick task in which the monkey released the joystick when a single peripheral square stimulus reduced its luminance. Many caudate neurons exhibited response-choice activity only during the two-patch condition (green dots in Fig 5E, n=28/80, AROC different from chance level 0.5 for two patches only) and did not show any side preference during the dimming condition (green dots in Fig 5F, n=21/70). Another group of caudate neurons showed a dependence on the location of the visual stimulus evoking the joystick release, and this side preference was retained across the three different task conditions (Fig 5E, violet dots, n=18/80), and also during the dimming task (Fig 5F, violet dots, n=21/70). For almost all of these neurons the side preference remained the same across visual conditions (Fig 5E, n=17/18) and tasks (Fig 5F, n=19/21).

Thus, like the visual cue-related modulation of caudate neurons described earlier, the response-choice activity was also often specific to the visual and task conditions – in particular, the presence of a visual distracter – even though the act of releasing the joystick was the same and was largely unaffected by which hand was used.

### Classification of epochs of the attention task based on caudate activity

Because caudate neurons exhibited diverse patterns of activity that appeared to cover the different trial conditions and the full timecourse of the attention task, we speculated that it might be possible to accurately identify the task epoch based on their activity alone. We used a linear classifier (support vector machine, SVM) to test whether there was sufficient information contained in the spike counts from our population of caudate neurons to correctly identify the epoch and cue condition in the attention task. As described in more detail in Material and methods, we identified 14 unique epochs based on the cueing condition and trial events in the attention task, trained a classifier for each of the epochs, and then cross-validated and quantified the performance of each classifier by testing it with spike counts from its own epoch as well as each of the 13 other epochs. To visualize the results, in Fig 6A we display the performance of each classifier for each of the 14 trial epochs in matrix format, using a color scale to indicate the percentage of test trials identified.

**Fig 6.**
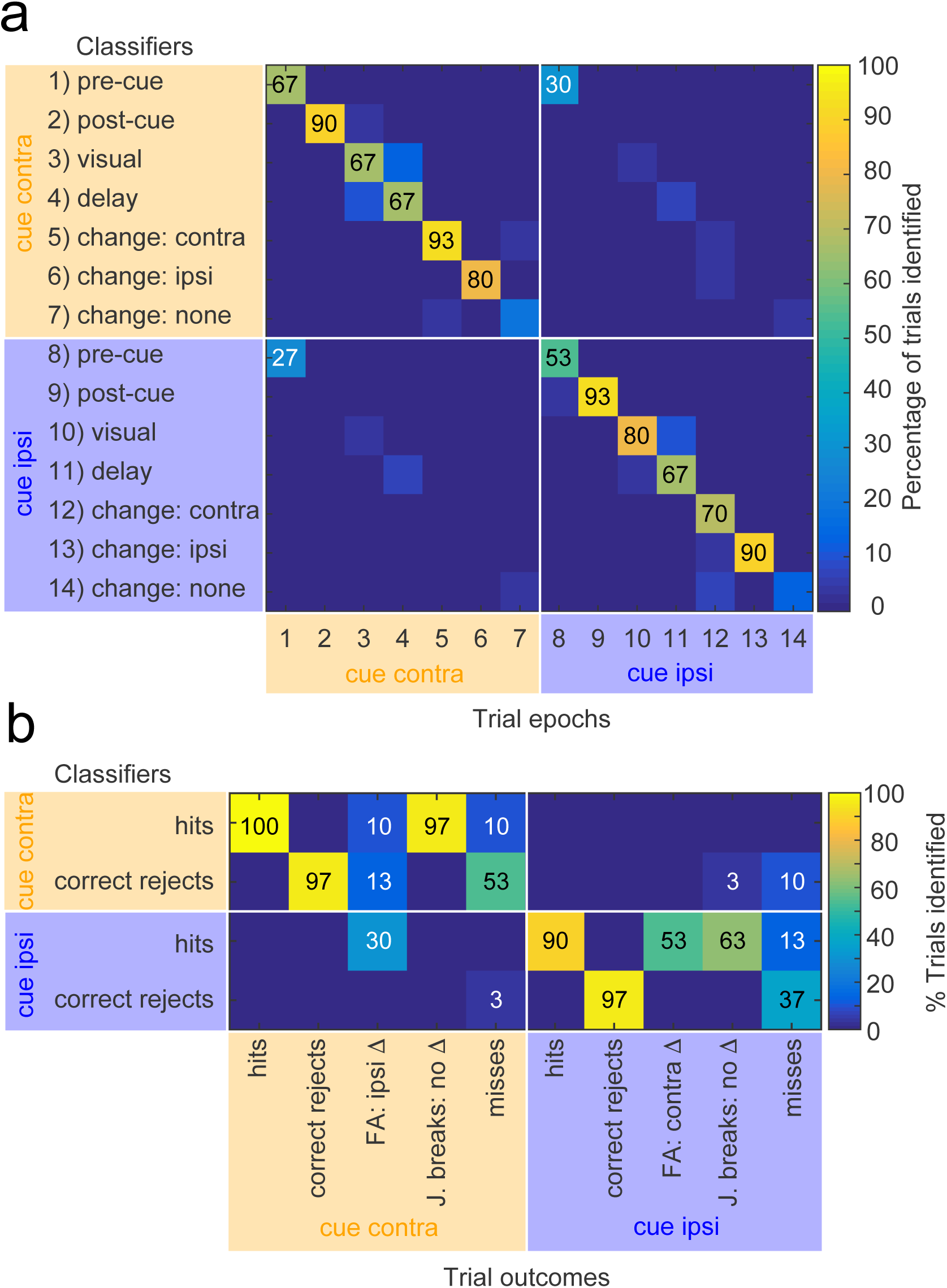
Linear classifier performance for the covert attention task. (A) Linear classifiers applied to caudate activity from different epochs of the attention task. The matrix illustrates the percentage of trials positively identified, as indicated by the color scale, by each of the 14 classifiers (rows) for each of the 14 trial epochs (columns). The diagonal of the matrix corresponds to the percentage of trial epochs correctly identified by each classifier; values outside the diagonal indicate the percentages of trial epochs that were incorrectly identified by classifiers trained on other epochs. Numbers overlaid on matrix indicate percentage scores for every case that was significantly greater than chance performance, for correct identifications (black numbers along diagonal) and for incorrect identifications (white numbers). (B) Linear classifier performance for trial outcomes during the covert attention task. The matrix illustrates the percentage of trials positively identified, as indicated by the color scale, by each of the 4 classifiers trained on change-epoch data from correct trials (rows) for each of the 10 possible trial outcomes (columns). Numbers overlaid on matrix indicate percentage scores for every case that was non-zero, for values greater than chance performance (black numbers) and for values less than chance performance (white numbers).

The set of classifiers performed significantly above chance for 12 of the 14 epochs, with no significant misclassifications except for the pre-cue epochs (Fig 6A). The post-cue epochs were classified with high accuracy (90 and 93% correct), regardless of whether the cue was contralateral or ipsilateral. The motion-change epochs were also classified accurately, especially when the change occurred at the cued location (e.g., 93% for contra change with contra cue, 90% for ipsi change with ipsi cue), with somewhat lower accuracy when the change occurred at the uncued location (80 and 70% for contra and ipsi respectively). The pre-cue epochs were also classified better than chance (67 and 53% correct), presumably reflecting the fact that cue conditions were blocked, although the two pre-cue epochs were also the only cases with significant, mutual misclassifications (30 and 27% errors), maybe because these epochs are less valuable of representing the task sequence compared to the other ones. The two no-change epochs (“change: none” in Fig 6A) were the only epochs that were not correctly identified above chance level.

These results quantitatively demonstrate the point suggested qualitatively in Fig 2B by the progression of peak activity across caudate neurons during the task sequence – there was sufficient information in the pattern of spike counts across our population of caudate neurons to identify with high probability almost all of the epochs of the attention task, during both contralateral and ipsilateral cueing conditions.

### Classification of trial outcomes during the attention task

Because caudate neuronal activity was related to response choice, and also contained sufficient information to identify epochs of the attention task (Fig 6A), we tested whether the spike counts from our population of caudate neurons could identify the trial outcomes during the attention task. Specifically, we sought to determine if caudate activity during the motion-change epoch was related to the stimulus condition or to the perceptual choice. To this end, we subdivided the data from the change epoch used previously for the linear classifier analysis (epoch 5-7 and 12-14, Fig 6A) based on the trial outcome (hit, correct reject, miss, false alarm and joystick break). Consistent with the previous analysis, we subdivided erroneous releases of joystick into two types – joystick releases for motion change on the uncued side (false alarms) and joystick releases for no motion change (joystick breaks). We adopted the strategy of training classifiers on data from the four types of correct trials (hits and correct rejects for contralateral and ipsilateral cue conditions), and then testing these four classifiers with data from all 10 trial outcomes.

The results demonstrate that caudate neuronal activity represents the response choice, correct or incorrect, especially when the cue is on the contralateral side (Fig 6B). Classifiers identified correctly performed trials (hits and correct rejects) with high probability (90-100%); these same classifiers also identified particular classes of errors. For example, the classifier for contralateral hits also identified 97% of joystick breaks committed during contralateral cues with no motion change (upper-left row of matrix in Fig 6B), consistent with erroneous detection of a motion change that did not happen. On the other hand, the contralateral hits classifier only rarely identified false alarms committed during contralateral cues with ipsilateral motion change (10%); these were more frequently identified by the ipsilateral hits classifier (30%), consistent with erroneous coding of the cue condition. The classifier for contralateral correct rejects also identified 53% of misses committed during contralateral cues, suggesting that the motion-direction change was simply not detected. For the ipsilateral classifiers, the outcomes were slightly different. The classifier for ipsilateral correct rejects also identified misses (37%), but the classifier for ipsilateral hits identified both types of erroneous releases (false alarms and joystick breaks) committed during ipsilateral cues (53% and 63%), albeit with lower probabilities.

These results quantitatively demonstrate that the pattern of spike counts across our caudate neurons represented the perceptual choices made during the covert attention task, on both correct and error trials. Choices during contralateral and ipsilateral cue conditions are both represented, but the patterns associated with erroneous releases (false alarms and joystick breaks) are different between the two cue conditions.

## Discussion

By recording neuronal activity in the caudate nucleus during a covert visual attention task, our results provide new insight into the role of the caudate in selection and covert attention processes. Because our task involved a non-spatial response (holding or releasing a joystick) guided by stimuli presented in the right or left visual field, we were able to document the visual-cue selectivity of caudate neurons during the covert attention task, dissociated from selectivity for the spatial goal of the movement. We also unexpectedly found that there was sufficient information in the pattern of activity across our population of caudate neurons to correctly identify the sequence of epochs during the visual attention task, as well as the response choice, regardless of cueing condition. These results are consistent with the view that the basal ganglia are important for representing the sequence of belief states that underlie action selection (Samejima and Doya 2007), and illustrate how this type of mechanism might apply to perceptual and cognitive functions even when no overt action is involved.

### Spatial selectivity of caudate neurons during covert attention task

We found that the activity of caudate neurons was strongly modulated by spatial cues during the covert attention task, consistent with previous findings that neurons in the caudate head are modulated by task instructions (Hikosaka et al. 1989; Apicella et al. 1992). Indeed, the size of the cue-related modulation we found for caudate neurons was much larger than what has been typically reported for visual cortical areas during visual attention tasks similar to ours. The firing rates of our caudate neurons were modulated by spatial cues by more than 100% (median modulation indices were 0.32-0.49, Fig 2), compared with the more modest changes of ∼8% found for V1 (McAdams and Maunsell 1999), 10-20% for MT (Seidemann et al. 1999), ∼26% for area V4 (McAdams and Maunsell 1999), ∼40% for MST (Treue and Maunsell 1996; 1999), and 25-50% found for area LIP (Herrington and Assad 2009; 2010). The attention cue-related changes we observed for caudate neurons are more consistent with the larger cueing effects found for neurons in frontal cortical areas such as the frontal eye fields (Armstrong et al. 2009) and dorsolateral prefrontal cortex (Lennert and Martinez-Trujillo 2013). Our caudate recordings also included a substantial number of neurons that showed higher activity for ipsilateral spatial cues, similar to what has been found for dorsolateral prefrontal cortex (Lennert and Martinez-Trujillo 2013), and unlike the preference for contralateral spatial cues found in visual cortex. These similarities in spatial cueing effects between frontal cortex and our caudate neurons are consistent with the known anatomy. The frontal cortex provides direct projections to regions in the head of the caudate nucleus that overlap with the locations of our recording sites (Ferry et al. 2000); visual cortical areas implicated in our motion-change task would be expected to project to more posterior regions around the genu of the caudate nucleus (Saint Cyr et al. 1990).

The large and bilateral spatial cueing effects we found for caudate neurons are presumably related to aspects of selective attention that lie downstream of changes in sensory processing. One possibility is that these cueing effects are related to spatial working memory. Selective attention and working memory are often studied as separate behavioral phenomena, but performing an attention task requires remembering the location of the cue, and conversely, there is substantial evidence that working memory may rely on rehearsal using visual selection mechanisms (Awh and Jonides 2001; Theeuwes et al. 2009). Neurons in dorsolateral prefrontal cortex are well known for the sustained delay-period activity that could support working memory (Funahashi et al. 1989; 1993), but this sustained activity is also consistent with representing the currently attended location (Lebedev et al. 2004). Our results demonstrate that similar signals are present on caudate neurons, and the design of our selective attention task allows us to attribute this activity to visual working memory rather than to movement planning because our task involved a non-spatial joystick response and no targeting saccades.

Another possibility is that the spatial selectivity was related to reward expectation, which is often closely linked to spatial attention (Maunsell 2004). For example, a previous study showed that caudate neurons can exhibit selective activity for visual cues that predict the direction of rewarded saccadic eye movements (Lauwereyns et al. 2002). In our task, the delivery of reward was not related to a goal-directed movement, because animals responded by releasing or holding down the same joystick across all task conditions. Thus, we can rule out the possibility that the spatial selectivity we observed was due to reward expectation for a spatially directed movement; instead, the spatial selectivity we observed was related to the location of the visual stimuli.

We also showed that the cue-related modulation of caudate neurons was strongly dependent on the visual and task conditions. One possible explanation is that this selectivity is a byproduct of movements that are planned but then not executed: subjects in our covert attention task might have preferred to look directly at the cued motion patch if they had been allowed to break central fixation. However, the properties of our caudate neurons are not consistent with this explanation. We found that the spatial selectivity exhibited during the attention task was not well predicted by the spatial preferences during memory guided saccades or the other joystick mapping task, indicating that it is unlikely that the cue-related modulation during the attention task was simply due to movement planning.

In addition, the visual-evoked activity of caudate neurons was specifically modulated during the covert attention task when the second, distracter patch was present, but not when only a single patch was presented. Similarly, the response-choice activity of many caudate neurons was specific to motion-direction changes in one visual hemifield, only when a distracter was present, even though the physical act of releasing the joystick was the same across all tasks, and did not involve a goal-directed movement. These results demonstrate that choice-related activity in the caudate includes the covert spatial selection of behaviorally relevant events, in addition to the overt preparation of orienting movements (Ding and Gold 2010; 2013).

We also found that in caudate nucleus, sensitivity and detect probability were correlated suggesting that the perceptual decision might be based on the activity of the most sensitive neurons; similar results have been shown in cortical areas MT (Masse et al. 2012) and LIP (Law and Gold 2008). These observations are consistent with the caudate being part of the circuit for implementing the perceptual decision during the covert attention task.

### Caudate activity and task sequencing

The cue-related modulation of our caudate neurons differed in another important way from what has been typically found in cortical areas during attention tasks. Rather than exhibiting a relatively constant cue-related preference during the task, we found that different subsets of caudate neurons showed phasic cue-related modulation during particular time epochs, so that the population of neurons tiled the full duration of the task, even during the response choice. Previous studies have demonstrated that caudate neurons can represent a task sequence when animals perform a sequence of oriented movements (Miyachi et al. 2002; Seo et al. 2012) and that caudate activity tiles the full duration of the trials (Lau and Glimcher 2007). Our results are substantively different from these previous findings, because our covert attention task did not include overt movements to the attended location, and thus demonstrate activity related to task sequence rather than action sequence. This type of sequential transient activation of caudate neurons is strikingly similar to a pattern recently reported in the dorsomedial striatum of rats during a non-match to position task (Akhlaghpour et al. 2016); as in our results, this sequential activity could not be explained by changes in motor activity, and was present over the entire task, not just the delay period. One might argue that the sequential activation of caudate neurons was simply related to time and not the task sequence itself. However, the cue-related modulation provides compelling evidence that activity depended on the sequence of events – for example, many caudate neurons showed visual- and choice-related activity that was specific to the visual field location of the preceding spatial cue even though the timing of events was the same across these conditions.

Our results also support the idea that the striatum plays a central role in representing the sequence of “belief states” needed to guide action selection (Samejima and Doya 2007). Using a linear classifier, we found that the pattern of activity across our population of caudate neurons correctly identified trial epochs throughout the visual attention task. Because most of these trial epochs involved maintained fixation and the absence of any overt action, these results demonstrate that caudate activity is not only important for tracking sequences of actions, as previously shown (Miyachi et al. 2002; Seo et al. 2012), but also for tracking the sequence of seemingly quiescent behavioral states that are necessary to perform a covert attention task (Krauzlis et al. 2014). During the motion-change epoch, the patterns of activity matched the perceptual choice regardless of whether it was correct or not (Fig 6b), consistent with caudate neurons representing the belief state of the subject.

This perspective nicely dovetails with recent evidence showing that caudate neuronal activity encodes several aspects of perceptual decision-making during a visual-motion direction-discrimination task (Ding and Gold 2010). As in our data set, the caudate neurons recorded by Ding and Gold (2010) showed a strong dependence on sensory and task conditions, a diversity of activity patterns and timing, and only a subset of neurons had activity linked to the action – in their case, a saccade to one of two choice targets. They interpret their results within the framework of accumulation of sensory evidence toward a decision boundary (Gold and Shadlen 2007), which has been an enormously fruitful approach; it may be useful now to consider how this sensory-based decision fits into the longer sequence of behavioral steps needed to perform the task. If it is true that the pattern of caudate activity represents a sequence of belief states, then each transition from one distinct pattern to another may correspond to a “decision”, even if the design of the experiment is mostly concerned with the particular state transition that is linked to the animal’s overt choice. This interpretation provides an answer to a long-standing question about the accumulator model, namely, what determines when the accumulation process should start. From the viewpoint of the sequence of behavioral states, the accumulation process would start with the “pre-choice” state that immediately precedes the “choice” state. The transition to this “pre-choice” state, presumably guided by other sensory events and the learned temporal structure of the task, is itself a type of decision, albeit a covert one. For example, in our case we observed ramp-like activity similar to that of Ding and Gold (2010), but it began during the delay period prior to the abrupt motion-direction change (Fig 2), suggesting that this was not an accumulation of sensory evidence but a gradual change in belief based on elapsed time (Jin et al. 2009), urgency (Carland et al. 2016) or other information.

### Conclusions

Our results provide new insights into the functions of the basal ganglia and how they may contribute to selective visual attention. We found that neurons in the primate caudate nucleus strongly select the spatial location of behaviorally relevant stimulus, even in the absence of any overt motor response, and that caudate neuron activity is sufficient to represent the sequence of events during the performance of a covert attention task. These results demonstrate that the role of the basal ganglia in selection and sequencing extends to non-motor aspects of behavior, and illustrate how the tracking of behavioral states by striatum and related circuits could support non-motor, cognitive functions such as attention and decision-making, as well as motor functions such as action selection.

**Author Contributions**
F.A. and R.J.K. designed the experiments, analysed the data, and wrote the manuscript. F.A. conducted the experiments.

## Acknowledgments

The authors thank Drs O. Hikosaka, B. Averbeck, L. Wang, and L. Katz for feedback on the manuscript. This work was supported by the National Eye Institute Intramural Research Program at the National Institutes of Health.

